# The landscape of *SNCA* transcripts across synucleinopathies: New insights from long reads sequencing analysis

**DOI:** 10.1101/524827

**Authors:** Elizabeth Tseng, William J Rowell, Omolara-Chinue Glenn, Ting Hon, Julio Barrera, Steve Kujawa, Ornit Chiba-Falek

**Author notes:** To whom correspondence should be addressed: Ornit Chiba-Falek Dept of Neurology DUMC Box 2900 Duke University, Durham, North Carolina 27710, USA.

## Abstract

Dysregulation of alpha-synuclein expression has been implicated in the pathogenesis of synucleinopathies, in particular Parkinson’s Disease (PD) and Dementia with Lewy bodies (DLB). Previous studies have shown that the alternatively spliced isoforms of the SNCA gene are differentially expressed in different parts of the brain for PD and DLB patients. Similarly, SNCA isoforms with skipped exons can have a functional impact on the protein domains. The large intronic region of the SNCA gene was also shown to harbor structural variants that affect transcriptional levels. Here we apply the first study of using long read sequencing with targeted capture of both the gDNA and cDNA of the SNCA gene in brain tissues of PD, DLB, and control samples using the PacBio Sequel system. The targeted full-length cDNA (Iso-Seq) data confirmed complex usage of known alternative start sites and variable 3’ UTR lengths, as well as novel 5’ starts and 3’ ends not previously described. The targeted gDNA data allowed phasing of up to 81% of the ~114kb SNCA region, with the longest phased block excedding 54 kb. We demonstrate that long gDNA and cDNA reads have the potential to reveal long-range information not previously accessible using traditional sequencing methods. This approach has a potential impact in studying disease risk genes such as SNCA, providing new insights into the genetic etiologies, including perturbations to the landscape the gene transcripts, of human complex diseases such as synucleinopathies.

## INTRODUCTION

Transcriptional and posttranscriptional programs control gene expression levels and/or production of multiple distinct mRNA isoforms, and changes in these mechanisms result in dysregulation of gene expression and differential expression profiles. Aberrant transcriptional and posttranscriptional gene regulation are abundant in human nervous system tissues and contributes to phenotypic differences within and between individuals in health and disease.

Dysregulation of alpha-synuclein expression has been implicated in the pathogenesis of synucleinopathies, in particular Parkinson’s Disease (PD) and Dementia with Lewy bodies and (DLB). While the role of *SNCA* overexpression in synucleinopathies, mainly PD, has been well established, here we focused on determination of the complete repertoire of *SNCA* transcript isoforms in different synucleinopathies. Previously, several different *SNCA* transcript isoforms have been described for *SNCA* gene, arisen from alternative splicing, transcriptional start sites (TSSs) and selection of polyadenylation sites (Xu, Tan, and Yu 2014; McLean et al. 2012). Alternative splicing of the coding exons gives rise to *SNCA* 140, *SNCA* 112, *SNCA* 126, and *SNCA* 98, resulting in four protein isoforms (Beyer and Ariza 2012). Alternative TSSs of *SNCA* gene results in four different 5’UTRs, and alternative selection of different polyadenylation sites determines three major lengths of the 3’UTR, with no impact on the composition of the protein product (Beyer and Ariza 2012). Our overarching goal is to gain new insights into the contribution of the different *SNCA* mRNA species, known and novel, to the pathogenesis and heterogeneity of synucleinopathies.

To date, most studies have used short read sequencing technologies to interrogate the transcriptome complexity in human brains. The availability of third generation long read technologies provides an unprecedented and nearly complete picture of isoform structures. However, existing long read transcript sequencing for human disease genes have used an amplicon-based approach (Treutlein et al. 2014; Tseng et al. 2017; Kohli 2017). While this approach has been successful in identifying complex alternative splicing in human disease genes, it is limited to the PCR primer design and will not uncover alternative start and end sites. Targeted enrichment, such as through the use of IDT probes, can deliver comprehensive isoform view of genes of interest at low sequencing cost. Further, highly accurate full-length transcript reads enable isoform-specific haplotyping.

Here, we present the first known study using targeted capture of gDNA and cDNA of the SNCA gene region using PacBio SMRT Sequencing. The SNCA gene region is ~114kb long, consisting of six exons with transcript lengths around 3kb. We multiplexed 12 human brain samples from PD, DLB, and Normal control samples, and sequenced the gDNA and cDNA library on the PacBio Sequel system. We describe the bioinformatics analyses used to identify SNPs, indels, and short tandem repeats for the gDNA capture, and isoform-level haplotyping for the cDNA data. We show that targeted capture is a cost-effective way of jointly studying genomic variation an alternative splicing in a disease-related neural gene.

## RESULTS

We designed custom probes for the SNCA gene and performed targeted capture of both gDNA and cDNA on a multiplexed library consisting of 12 human brain samples from PD, DLB, and normal controls (Figure 1, Supplementary Table 1). The gDNA and cDNA libraries were sequenced on the PacBio Sequel platform. Bioinformatics analysis was done using PacBio software followed by custom analysis.

**Figure 1.**
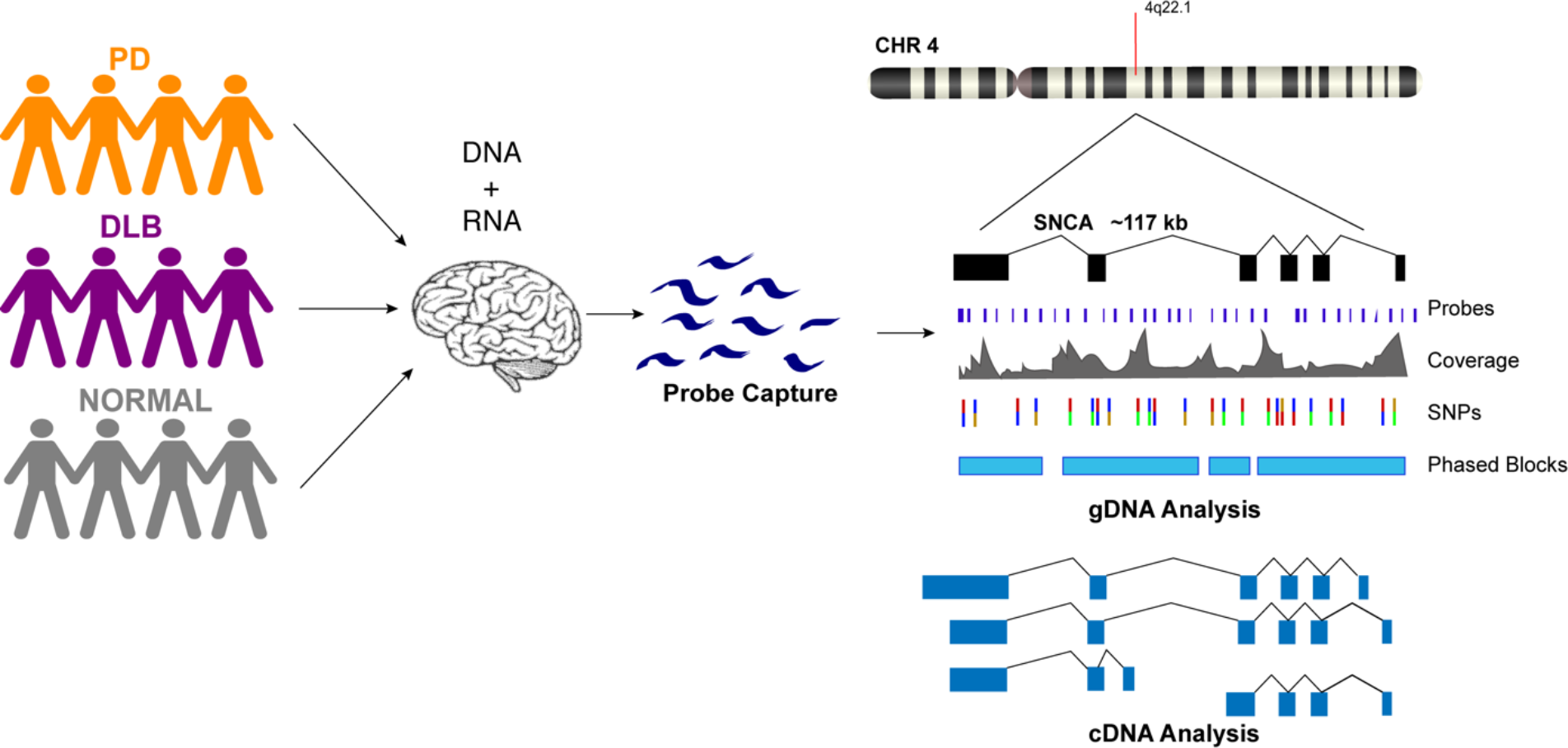
Schematic presentation of the study design. DNA and RNA material were extracted from postmortem brain tissues of patients from Parkinson’s disease, Dementia with Lewy Body, and control groups. gDNA and cDNA libraries were made using probe hybridization and sequenced on the PacBio Sequel platform. Analysis was performed using PacBio software and other existing tools.

### Targeted gDNA capture identified known and novel variations

After generating circular consensus sequences (CCS) and removing PCR duplicates (Supplemental Methods), we obtained 16-to 71-fold mean unique coverage of the SNCA gene region. The CCS reads had a mean insert length of 2.9 kbp and a mean read accuracy of 98.9%. With the exception of a 5 kbp region intentionally uncovered by probes due to the presence of LINE elements (hg19 chr4:90,697,216-90,702,113) and a 2.1 kbp region of high GC content around exon 1, there was sufficient coverage to genotype both haplotypes for each of the 212 samples (Figure 2, Supplementary Figure 1).

**Figure 2.**
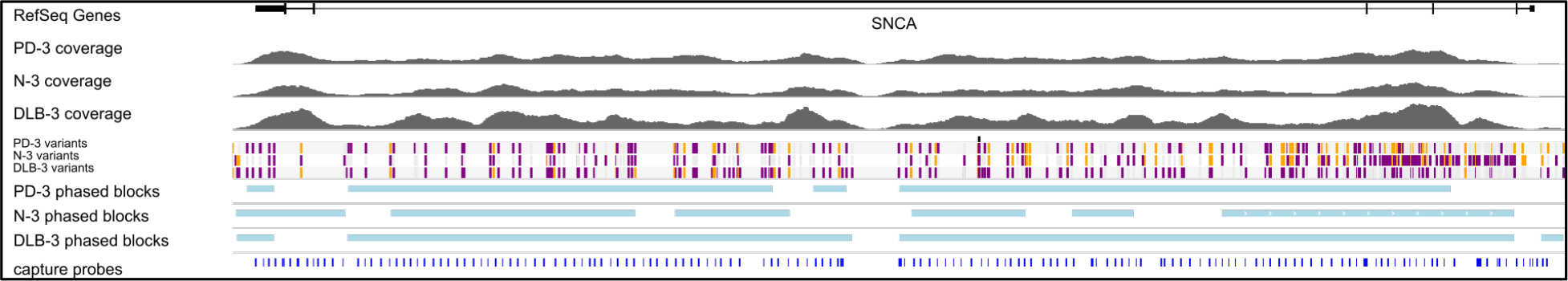
Targeted gDNA capture and phasing. An example showing one sample from each condition. Top track shows one of the SNCA isoforms, followed by the gDNA coverage for the three samples. The variant track shows each SNP and are color-coded for heterozygous (purple), homozygous alternative (orange), and homozygous reference (gray). Phased blocks are shown in light blue. Bottom track shows capture probe locations. The dropout region in probe design are due to two LINE elements in the middle of intron 4. For the gDNA coverage and phasing information of all 12 samples, see Supplementary Figures.

Using GATK HaplotypeCaller (GATK HC), quality-based filtering, and manual curation, we identified 282 SNPs and 35 indels, including novel 8 SNPS and 13 indels not found in dbSNP (Supplementary Table 2). No variants were identified in the coding region for SNCA, although 8 variants were identified in untranslated regions. The majority of the identified variants, including several short tandem repeats (STR), fall within introns 2, 3, and 4.

We have previously described a highly polymorphic CT-rich region in intron 4 of SNCA with 4 observed haplotypes (Lutz et al. 2015). While this highly repetitive and structurally variable region proved difficult to genotype with GATK4 HC, we were able to construct consensus sequences for all 12 samples and observed all 4 of the previously discovered haplotypes (Supplementary Figure 2). Additionally, we identified a novel STR in intron 4 consisting of a three-base unit repeated 16 times in the reference. Within the twelve samples, we identified three haplotypes, with 9, 12, and 15 copies of the TTG repeat unit. GATK HC correctly genotyped all of these except for one haplotype for PD-4, which had fairly low coverage in this region. However, with the given data for this sample, the genotype can be determined by visual inspection (Table 1).

**Table 1.** A novel triplet tandem repeat in intron 4. (chr4: 90713442, (TTG)16 reference). The reference has 16 repeats. The table shows the repeat number of both haplotypes for each sample. *PD-4 is incorrectly genotyped by GATK4HC but can be genotyped by visual inspection.

| Sample | Genotype |
| --- | --- |
| PD-1 | (TTG)<sub>15</sub> / (TTG)<sub>15</sub> |
| PD-2 | (TTG)<sub>12</sub> / (TTG)<sub>12</sub> |
| PD-3 | (TTG)<sub>12</sub> / (TTG)<sub>12</sub> |
| PD-4 | (TTG)<sub>15</sub>* / (TTG)<sub>12</sub> |
| N-1 | (TTG)<sub>15</sub> / (TTG)<sub>12</sub> |
| N-2 | (TTG)<sub>12</sub> / (TTG)<sub>12</sub> |
| N-3 | (TTG)<sub>12</sub> / (TTG)<sub>12</sub> |
| N-4 | (TTG)<sub>12</sub> / (TTG)<sub>12</sub> |
| DLB-1 | (TTG)<sub>12</sub> / (TTG)<sub>12</sub> |
| DLB-2 | (TTG)<sub>12</sub> / (TTG)<sub>12</sub> |
| DLB-3 | (TTG)<sub>15</sub> / (TTG)<sub>9</sub> |
| DLB-4 | (TTG)<sub>12</sub> / (TTG)<sub>12</sub> |

We used the short variants detected by GATK HC in conjunction with the read-based phasing tool WhatsHap (Martin et al. 2016) to phase the CCS reads across the locus, with a range of success driven mostly by the heterozygous variant density over the locus. Samples PD-1, PD-4, N-4, DLB-1, and DLB-4 had long stretches of low heterozygosity, with very few, short phase blocks, while the other samples yielded phase blocks ranging from 7 to 18 times the mean read length, up to 54 kbp (Supplementary Figure 3).

### Targeted cDNA capture identified novel start and end sites

We processed the PacBio cDNA (Iso-Seq) data using the PacBio SMRTAnalysis software. After mapping the Iso-Seq data to hg19 and removing artifacts (Supplementary Table 3, Supplementary Figure 4), we obtained a final set of 41 SNCA isoforms (Figure 3). All final isoforms have all canonical splice sites and are supported by a total of 20 or more full-length reads. The majority of the isoforms (28 out of 41) have all six exons, with differences coming solely from the use of alternative 5’ start sites and different 3’ UTR lengths. The 3’ UTR lengths varied between 300 bp to 2.6 kb. The use of highly diverse alternative 5’ start site in SNCA is known; what is less known is the variable 3’ UTR length, which had been previously studied using RNA-seq data that did not resolve full-length isoform structures (Rhinn et al. 2012). The Iso-Seq data shows that the variable 3’ UTR length seems paired with all possible combinations of 5’ start sites and that there is no preferential coupling. Almost none of the variability in start and end site changes the predicted open reading frame (Supplementary Figure 5) and is predicted to translate to the canonical 141 amino acid sequence.

We observed two isoforms with exon 3 skipping (SNCA126) and five isoforms with exon 5 skipping (SNCA112). Again, the splicing diversity in these two exon skipping groups mostly came from the diverse use of alternative 5’ start sites and variable 3’ UTR length. ORF prediction shows skipping exon 3 or exon 5 shortens the ORF but maintains the reading frame. Three isoforms have novel 3’ end sites located in intron 4. ORF prediction shows that this results in truncated protein product.

We identified a previously unannotated 5’ start site located in intron 4 (hg19 coordinate chr4: 90692548-90693045, Figure 3). The three isoforms associated with this novel start consists of the novel start site, exon 5, and variable 3’ UTR lengths. Interestingly, while publicly downloaded short read data from GTEx and Sandor et al.(Sandor et al. 2017) did not support this novel start site, a recent public NA12878 direct RNA dataset (https://github.com/nanopore-wgs-consortium/NA12878/) contained only one SNCA transcript that confirmed this alternative start site. Interestingly, this novel 5’ start site is predicted to introduce novel peptides while maintaining the reading frame in exon 5.

**Figure 3.**
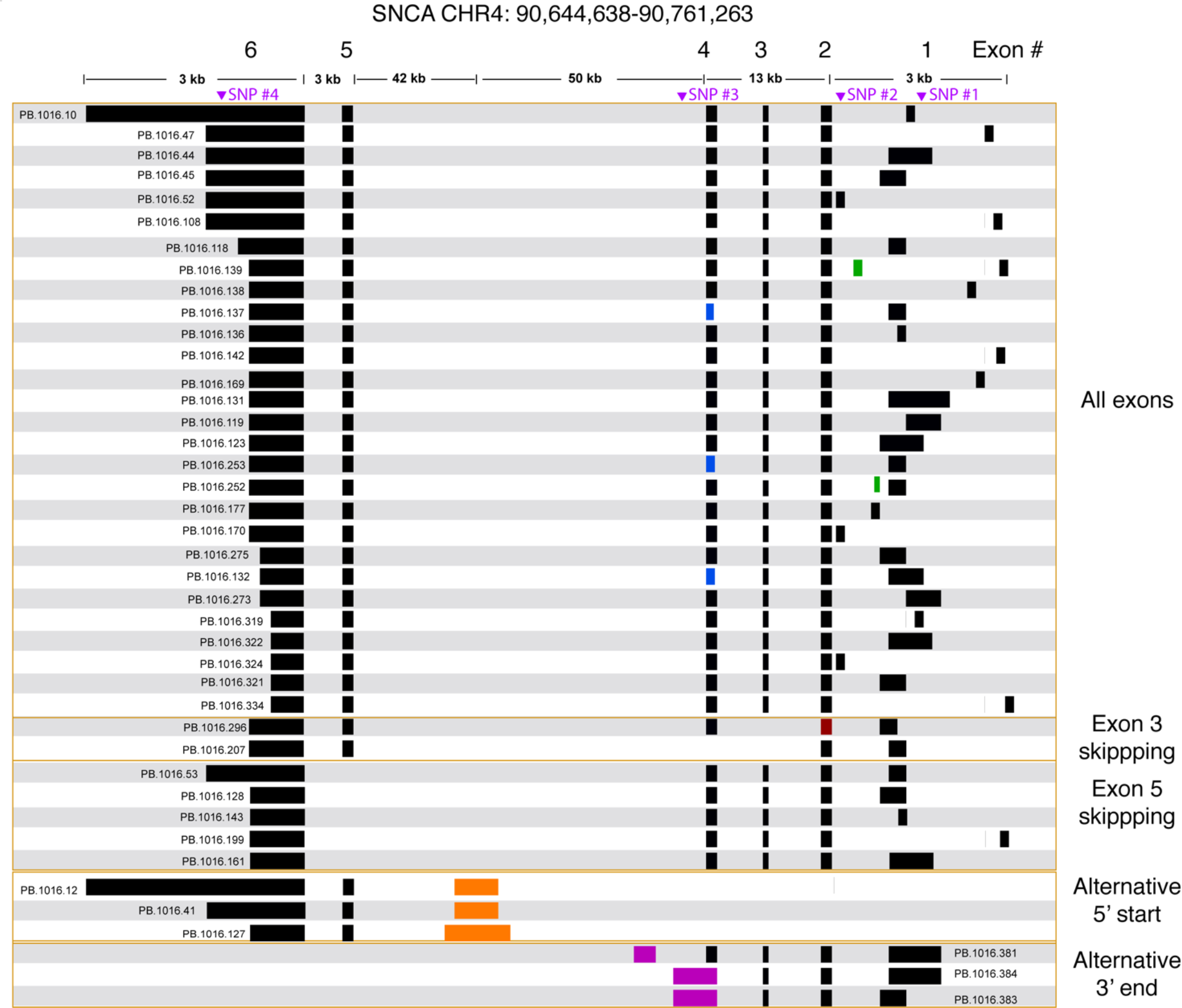
SNCA isoforms captured using targeted Iso-Seq identifies novel start and end sites. (a) Final set of 41 SNCA isoforms. The majority of the isoform complexity comes from combinatorial usage of alternate 3’ UTR lengths and exon 1, with a few rare alternative splice sites found in exon 1 (green), 2 (red), and 4 (blue). All junctions have canonical splice sites. We identified five isoforms that skipped exon 5 and two isoforms that skipped exon 3. We also identified novel start (orange) and end sites (purple) in intron 4. Called SNPs are marked in purple.

We also identified three SNCA transcripts with new end sites (Figure 3). Two isoforms (PB.1016.383, PB.1016.384) used an extended 3’ UTR in exon 4, while the third isoform (PB.1016.381) used a novel 3’ exon in intron 4. The novel 3’ UTRs results in a truncated ORF prediction.

Using the normalized full-length read count as a proxy for isoform abundance, we find one of the canonical SNCA isoforms (PB.1016.131) to be the most abundant, with an abundance of 50-60% across all subject samples (Supplementary Table 4). We further grouped the 41 isoforms by their splicing patterns (Table 2). Isoforms that have all 6 exons account for 95-97% of the abundance. Previous studies have shown a marked expression increase in isoforms missing exon 3 (SNCA126) in the frontal cortex of DLB samples compared to normal (Beyer et al. 2008); our aggregated isoform counts shows that three of the DLB samples have a slightly elevated count level compared to the normal samples as well as the SNCA112 (exon 5 skipping) variants for PD and DLB against normal samples.

**Table 2.** SNCA isoform abundance for each sample, aggregated by splicing patterns. The abundance for each isoform is the fraction of on-target, full-length reads associated with that isoform. Isoforms are grouped by their splice patterns (see Figure 3). *AllExons*: isoforms expressing all 6 exons; *Skip3*: isoforms skipping exon 3; *Skip5*: isoforms skipping exon 5; *Alt5*: isoforms with the alternative start site in intron 4; *Alt3*: isoforms with the alternative end site in intron 4. The full abundance for each isoform is shown in Supplementary Table and Supplementary Data.

| GROUP | PD-1 | PD-2 | PD-3 | PD-4 | N-1 | N-2 | N-3 | N-4 | DLB-1 | DLB-2 | DLB-3 | DLB-4 |
| --- | --- | --- | --- | --- | --- | --- | --- | --- | --- | --- | --- | --- |
| **AllExons** | 95.92% | 97.06% | 95.82% | 95.76% | 98.10% | 97.73% | 96.63% | 96.68% | 96.00% | 94.59% | 94.58% | 96.21% |
| **Skip3** | 0.00% | 0.00% | 0.03% | 0.08% | 0.03% | 0.09% | 0.02% | 0.25% | 0.08% | 0.13% | 0.24% | 0.00% |
| **Skip5** | 2.50% | 1.47% | 1.68% | 1.10% | 0.79% | 0.93% | 1.00% | 1.32% | 2.04% | 1.42% | 1.56% | 1.89% |
| **Alt5** | 1.02% | 0.00% | 1.61% | 2.76% | 0.64% | 0.77% | 1.85% | 0.08% | 0.94% | 2.83% | 3.16% | 0.79% |
| **Alt3** | 0.56% | 1.47% | 0.85% | 0.30% | 0.44% | 0.49% | 0.51% | 1.67% | 0.94% | 1.03% | 0.47% | 1.10% |

### Full-Length cDNA enables isoform-level phasing information

We called SNPs using cDNA by piling up all full-length reads from the 12 samples to call variants (see Methods). A total of four SNPs were called and all were previously annotated in dbSNP (Table 3, Figure 3). The four SNPs are all located in non-CDS regions, one in the 3’ UTR (exon 6), one in intron 4, and two in the 5’ UTR (exon 1). The 3’ UTR SNP (chr4: 90646886) is only covered by isoforms with a 3’ UTR that is at least ~1kb long, and hence, not all canonical isoforms cover this SNP. The intron 4 SNP (chr4: 90743331) is only covered by the novel alternative 3’ end isoforms (PB.1016.383, PB.1016.384) and is not connected to any of the other SNPs. The two 5’ UTR SNPs (chr4: 90757312 and chr4: 90758389) are covered by two mutually exclusive exon 1 usage and hence, are also not linked.

Our current approach is limited to calling only substitution variants in transcribed regions with sufficient coverage. Comparing the list of our SNPs with the hg19 dbSNP annotation shows that most of the SNPs or variants missed were either less than 1% frequency in the population, were not single nucleotide substitutions, or adjacent to low complexity regions. For example, rs77964369 (chr4:90646532) is reported to have 50/50 frequency of T/A, however this T is adjacent to a stretch of 11 genomic As downstream. Manual inspection of the Iso-Seq read pileup, which has ~1300 reads at this site, does not suggest evidence of variation at least amongst our 12 samples.

Using the sample-specific reads, we call the genotype of each sample at each SNP location (Table 3). Besides PD-2 having too few reads and is inconclusive for all 4 SNPs, we were able to call the genotype for most other samples. Notably, DLB-3 was the only sample that is heterozygous at all SNP locations. Otherwise, we did not observe any condition-specific pattern of preferring one genotype to other.

**Table 3.** cDNA SNP information. SNPs were called using full-length reads from the Iso-Seq data. For each sample, the number of FL reads supporting either the reference or alternative base was tabulated. If both alleles have 5+ FL read support, it is called heterozygous; if one allele has 5+ reads and the other has < 5, homozygous; otherwise, inconclusive.

| # | Coord | dbSNP | Annotation | Ref | Alt | Homo_Ref | Homo_Alt | Het | Inconclusive |
| --- | --- | --- | --- | --- | --- | --- | --- | --- | --- |
| 1 | 90758389 | rs2301135 | exon1<br>(5' UTR) | G | C | PD-1,<br>DLB-1 | PD-3,<br>DLB-1,<br>DLB-2 | PD-4,<br>N-1,<br>N-2,<br>N-3,<br>DLB-3 | PD-2,<br>DLB-4 |
| 2 | 90757312 | rs2870027 | exon1<br>(5' UTR) | C | T | PD-3,<br>PD-4,<br>N-1,<br>N-2,<br>N-3,<br>N-4,<br>DLB-1,<br>DLB-2 |  | DLB-3 | PD-1,<br>PD-2,<br>DLB-4 |
| 3 | 90743331 | rs10005233 | intron4<br>(alt 3' UTR) | C | T | PD-1,<br>PD-4,<br>N-4,<br>DLB-4 | PD-3,<br>DLB-1,<br>DLB-2 | N-1,<br>N-2,<br>N-3,<br>DLB-3 | PD-2 |
| 4 | 90646886 | rs356165 | exon6<br>(3' UTR) | G | A | N-4,<br>DLB-4 | PD-1,<br>DLB-1 | PD-3,<br>N-1,<br>N-2,<br>N-3,<br>DLB-2,<br>DLB-3 | PD-2 |

## DISCUSSION

We describe the first study using targeted enrichment of the *SNCA* gene on multiplexed gDNA and cDNA libraries for studying neurological diseases using long read sequencing. The long readlengths of the PacBio Sequel system faciliated the sequencing of the full-length transcript isoforms repertoire of the SNCA gene. We revealed the diversity in the use of alternative 5’ start sites and variable 3’ UTR lengths and observed known exon skipping events, such as exon 3 deletion (SNCA126) and exon 5 deletion (SNCA112). Additionally, new alternative start and end sites within the large intron 4 were identified that are predicted to be translated to novel proteins. Noteworthily, the novel transcript isoforms were niether described previously nor identified in the analysis of short read transcriptomic data sets, possibly because long reads encompass the full-length of the different transcripts and the capture vs whole genome approach allows a deeper sequencing coverage.

The biological and pathological significance of the different SNCA protein isoforms has yet to be fully discovered. However, specific SNCA post translational modification and splicing isoforms have been associated with intracellular aggregation propensities(Kalivendi et al. 2010) and are differently expressed in human synucleinopathies (Beyer and Ariza 2012; Beyer et al. 2008). Studies of SNCA post translational modification showed that Lewy bodies, the pathological hallmark of synucleinopathies, contain abundant phosphorylated, nitrated, and monoubiquitinated SNCA (Kim, gedal, and Halliday 2014). The effects of post-transcriptional modification on SNCA aggregation have also been studied. Alternative splicing was suggested to affect SNCA aggregation. A deletion of either exon 3 or 5 predicts functional consequences: while exon 3 deletion (*SNCA*126) leads to the interruption of the N-terminal protein-membrane interaction domain which may lead to less aggregation, exon 5 deletion (*SNCA*112) may result in enhanced aggregation due to a significant shortening of the unstructured C-terminus (H.-J. Lee, Choi, and Lee 2001; Beyer 2006). In the frontal cortex of DLB, *SNCA*112 is increased markedly compared to the controls (Beyer et al. 2008), while *SNCA*126 levels are decreased in the prefrontal cortex of DLB patients (Beyer et al. 2006). In contrast, *SNCA*126 expression showed increased in the frontal cortex of PD brains and no significant differences in MSA (Beyer et al. 2008). *SNCA*98 is a brain specific splice variant that lacks both exon 3 and 5, and exhibits different expression levels in various areas of fetal and adult brain. Overexpression of *SNCA* 98 has been reported in DLB, PD (Beyer et al. 2007) and MSA (Beyer et al. 2008) frontal cortices compared with controls. In addition, the post transcriptional process resulting in alternative 3’UTR usage was reported to have effects on the mRNA stability and localization (Fabian, Sonenberg, and Filipowicz 2010; Rhinn et al. 2012; Yeh and Yong 2016). While further investigations regarding of the aggregation propensities of the different known SNCA protein isoforms and the composition of Lewy bodies are warrant, here, we discovered novel transcripts that in future studies their predicted translation to proteins need to be validated followed by in depth characterization of their cellular localization, presence in Lewy body and tendency to form aggregates.

In this paper, we focused on creating a sequencing and analysis standard for analyzing targeted gDNA and cDNA data generated from the same subject. This is a powerful approach that potentially allows the phasing of the gDNA sequences across the complete region of a particular gene based on heterozygosity in the sequence ofthe full-length transcript isoforms. The PacBio targeted gDNA data in this study produced phased blocks that covered 81% of the 114kb region centered on SNCA, with the longest phased block excedding 54kb. Upon the feasibility to generate longer read length and higher accuracy, the gDNA phasing blocks will likely increase. Also, the ability to phase was limited by the closeness of SNPs and the absence of heterozygosity in the cDNA data.

gDNA variants analysis confirmed known and identified novel short tandem repeats (STRs) in the intronic regions. For example, previously using phased sequencing by cloning and Sanger sequencing, we discovered four distinct haplotypes within an intronic CT-rich region that comprised of a cluster of variable repetitive sequences (Lutz et al. 2015). We showed that a specific haplotype, termed haplotype 3, conferred risk to develop Lewy body pathology in Alzheimer’s patients. Here, we validated the sequence of this highly polymorphic low-complexity region and its four defined haplotypes. In addition, confined by our sample size is very small (8 chromosomes each pathological group), we confirmed that the Lewy body pathology risk haplotype 3 is present exclusively in disease, *i.e.* PD (1 chromosome) and DLB patients (2 chromosomes). The pilot results and our previous publication provide the premise to reapeat the association analyses of synucleinopathies with accurately defined, *i.e.* by long reads, STRs and structural haplotypes using a larger sample size.

Our paper demonstrated the ability of the PacBio Sequel system to discover novel full-length transcripts and characterize the complete full-length transcripts repertoire of a gene implicated in a disease, such as *SNCA* in synucleinopathies presented in this work. Furthermore, we also showed that long reads gDNA define more accuretly short structural variants and haplotypes including STRs and by that can facilitate the discovery and validation of disease associated variants other that SNPs. Collectively, this new knowledge is highly valuable and applicable in advancing our understanding of the genetic etiologies, that may involve perturbations in the transcripts landscape, underlying human complex diseases including, age related neurodegenerative disorders such as synucleinopathies.

## METHODS

### Study Samples

The study cohort (N=12) consisted of individuals with three autopsy-confirmed neuropathological diagnoses: (1) PD (N=4); (2) DLB (N=4); and (3) clinically and neuropathologically normal subjects (N=4). Frontal cortex brain tissues were obtained through the Kathleen Price Bryan Brain Bank (KPBBB) at Duke University, the Banner Sun Health Research Institute Brain and Body Donation Program (Beach et al. 2015), and Layton Aging & Alzheimer’s Disease Center at Oregon Health and Science University. Neuropathologic phenotypes were determined in *postmortem* examination following standard well-established methods following the method and clinical practice recommendations of McKeith and colleagues(McKeith, Perry, and Perry 1999; McKeith et al. 2005). The density of the LB pathology (in a standard set of brain regions) received scores of mild, moderate, severe and very severe. The neurologically healthy brain samples were obtained from *postmortem* tissues of clinically normal subjects who were examined, in most instances, within 1 year of death and were found to have no cognitive disorder or parkinsonism and neuropathological findings insufficient for diagnosing PD, Alzheimer’s disease (AD), or other neurodegenerative disorders. All samples were whites. Demographics and neuropathology for these subjects are summarized in Table 1. The project was approved by the Duke Institution Review Board (IRB). The methods were carried out *in accordance with* the relevant guidelines and regulations.

### Genomic DNA and RNA extractions

Genomic DNA was extracted from brain tissues by the standard Qiagen protocol (Qiagen, Valencia, CA). Total RNA was extracted from brain samples (100 mg) using TRIzol reagent (Invitrogen, Carlsbad, CA) followed by purification with a RNeasy kit (Qiagen, Valencia, CA), following the manufacturer’s protocol. gDNA and RNA concentration were determined spectrophotometrically and the quality of the RNA samples and lack of significant degradation was confirmed by measurements of the RNA Integrity Number (RIN, Table 1) utilizing an Agilent Bioanalyzer.

### Library Preparation and Sequencing

#### gDNA capture using IDT xGen^®^ Lockdown^®^ Probes and single-molecule sequencing

2µg of each gDNA sample was sheared to 6kb using the Covaris g-TUBE and ligated with barcoded adapters. An equimolar pool of 12-plex barcoded gDNA library (2µg total) was input into the probe based capture with a custom designed SNCA gene panel.

A SMRTBell library was constructed using 626ng of captured and re-amplified gDNA (https://www.pacb.com/wp-content/uploads/Procedure-Checklist-%E2%80%93-Multiplex-Genomic-DNA-Target-Capture-Using-IDT-xGen-Lockdown-Probes.pdf). A total of 3 SMRT Cells (6 hour movie) were sequenced on the PacBio Sequel platform using 2.0 chemistry.

#### cDNA capture using IDT xGen^®^ Lockdown^®^ Probes and single-molecule Isoform-Sequencing (Iso-Seq)

100-150ng of total RNA per reaction was reverse transcribed using the Clontech SMARTer cDNA synthesis kit and 12 sample specific barcoded oligo dT (with PacBio 16mer barcode sequences, see Supplementary Methods). Three reverse transcription (RT) reactions were processed in parallel for each sample. PCR optimization was used to determine the optimal amplification cycle number for the downstream large-scale PCR reactions. A single primer (primer IIA from the Clontech SMARTer kit 5’ AAG CAG TGG TAT CAA CGC AGA GTA C 3’) was used for all PCR reactions post-RT. Large scale PCR products were purified separately with 1X AMPure PB beads and the bioanalyzer was used for QC. An equimolar pool of 12-plex barcoded cDNA library (1µg total) was input into the probe based capture with a custom designed SNCA gene panel.

A SMRTBell library was constructed using 874ng of captured and re-amplified cDNA (https://www.pacb.com/wp-content/uploads/Procedure-Checklist-%E2%80%93-cDNA-Capture-Using-IDT-xGen-Lockdown-Probes.pdf). One SMRT Cell (6 hour movie) was sequenced on the PacBio Sequel platform using 2.0 chemistry.

### gDNA analysis

Sequencing of the barcoded gDNA data was run on three Sequel 1M cells using 2.0 chemistry. The data were demultiplexed by running the Demultiplex Barcodes application in PacBio SMRT Link v6.0.

#### Short variant analysis and phasing

Circular Consensus Sequence (CCS) reads were generated using SMRT Analysis 6.0.0 from each demultiplexed dataset and aligned to the hg19 reference genome using minimap2. PCR duplicates from post-capture amplification were identified by mapping endpoints and tagged using a custom script. Short variants were called using GATK4 (Poplin et al. 2018). After a first pass of filtering using coverage depth and quality metrics, variants were manually inspected in IGV (http://www.broadinstitute.org/igv). If variants did not phase with nearby SNPs, they were manually filtered. The variant sites that passed manual curation were used in conjunction with the deduplicated CCS alignments for read-backed phasing with WhatsHap (Martin et al. 2016).

#### Clustering and determining haplotypes for CT-rich region

Subsequences aligned to chr4:90742331-90742559 (hs37d5) were extracted for each sample. After inspecting the size distribution of these subsequences, they were clustered by size and sequence similarity using a combination of python and MUSCLE [ref], and a consensus sequence was generated independently for each cluster.

Custom scripts and workflows further described in Supplementary Methods and https://gist.github.com/williamrowell/dc4fed90098da8af795bb10a434cbb3c

### Isoform Analysis

Sequencing of the barcoded cDNA data was on one Sequel 1M cell using 2.0 chemistry. Bioinformatics analysis was done using the Iso-Seq 3 application in the PacBio SMRT Analysis v6.0.0 to obtain high-quality, full-length isoform sequences (see Supplementary Methods for more information).

#### Isoform SNP Calling

Full-length reads associated with the final 41 isoforms from all 12 samples were aligned to the hg19 genome to create a pileup. Bases with QV less than 13 were excluded. Then, at each position with at least 40 base coverage, a Fisher exact test with Bonferroni correction is applied with a p-value cutoff of 0.01. Only substitution SNPs not close to homopolymer regions (stretches of 4 or more of the same nucleotide) were called. After SNP calling, the genotype for each sample was determined by tallying the number of supporting sample-specific FL reads. If a sample had 5+ FL reads supporting both reference and alternative base, it was heterogyzous. If a sample had 5+ FL reads supporting one allele and fewer than 5 FL reads for the other, it was homozygous. Otherwise, it was inconclusive. Scripts are available at: https://github.com/Magdoll/cDNA_Cupcake

## DATA ACCESS

The three SMRT cells of gDNA raw data is available at Zenodo.org with doi: 10.5281/zenodo.1560688. The one SMRT cell of cDNA raw data is available at Zenodo.org with doi: 10.5281/zenodo.1581809. The processed gDNA and cDNA results, including gDNA variants and cDNA isoforms, are available at Zenodo.org with doi: 10.5281/zenodo.1588732.

